# AZD5069 Inhibits Angiogenesis Without Cytotoxicity In Human Endothelial Cell Culture

**DOI:** 10.64898/2026.06.12.731993

**Authors:** Carlo R Bartoli, Autumn Anthony, Rohini Desetty

## Abstract

**Background:** The CXCR2 receptor pathway plays a major role in inflammatory and invasive angiogenesis in human disease.

**Objective:** We evaluated AZD5069, a selective CXCR2 antagonist, as an angiogenesis inhibitor in human cell culture.

**Methods:** Human Umbilical Venous Endothelial Cells (HUVECs), Human Aortic Endothelial Cells (HAECs), and Human Pulmonary Artery Endothelial Cells (HPAECs) were cultured with standard *in vitro* techniques. AZD5069 (0, 8, 16, 32, 64, 128, 256 μM) was evaluated as an angiogenesis inhibitor with fluorescent-labeled 5-Ethynyl-2’-deoxyuridine (EdU) uptake to quantify endothelial cell proliferation, scratch assay to quantify endothelial cell migration, and Geltrex assay to quantify endothelial cell tubule and hub formation. AZD5069 cytotoxicity was evaluated with *in situ* terminal deoxynucleotidyl transferase 2’-Deoxyuridine triphosphate-‘5 (dUTP) nick-end labeling (TUNEL) to quantify apoptosis and membrane-impermeable cyanine dye uptake to quantify necrotic cell death.

**Results:** AZD5069 significantly reduced HUVEC, HAEC, and HPAEC proliferation, migration, tubule count, total tubule length, and node count with a dose-response. AZD5069 did not cause apoptosis nor necrotic cell death.

**Conclusions:** AZD5069 inhibited angiogenesis without cytotoxicity in human endothelial cell culture. The endothelial cell CXCR2 receptor pathway may be a novel target for anti-angiogenesis therapy. AZD5069 may have clinical utility in cardiovascular, oncologic, and inflammatory disease.

**Condensed Abstract:** The CXCR2 receptor pathway plays a major role regulating angiogenesis in inflammation and cancer. The CXCR2 receptor pathway has been evaluated in humans as a target for therapy in inflammatory disease and cancer but not as a therapeutic approach to block pathologic angiogenesis. AZD5069 is a clinical stage, direct CXCR2 antagonist. In human endothelial cell culture, AZD5069 inhibited angiogenesis without causing apoptosis or necrotic cell death. The endothelial cell CXCR2 receptor pathway may be a novel target for anti-angiogenesis therapy. AZD5069 may have clinical utility as a novel angiogenesis blocker in human disease.

Visual Abstract:
AZD5069, a selective CXCR2 antagonist, significantly reduced endothelial cell proliferation, migration, and vascular tubule formation without causing necrotic or apoptotic cell death. The endothelial cell CXCR2 receptor pathway may be a novel target for anti-angiogenesis therapy. AZD5069 may have clinical utility in human cardiovascular, oncologic, and inflammatory disease with pathologic, dysregulated, or excessive angiogenesis.

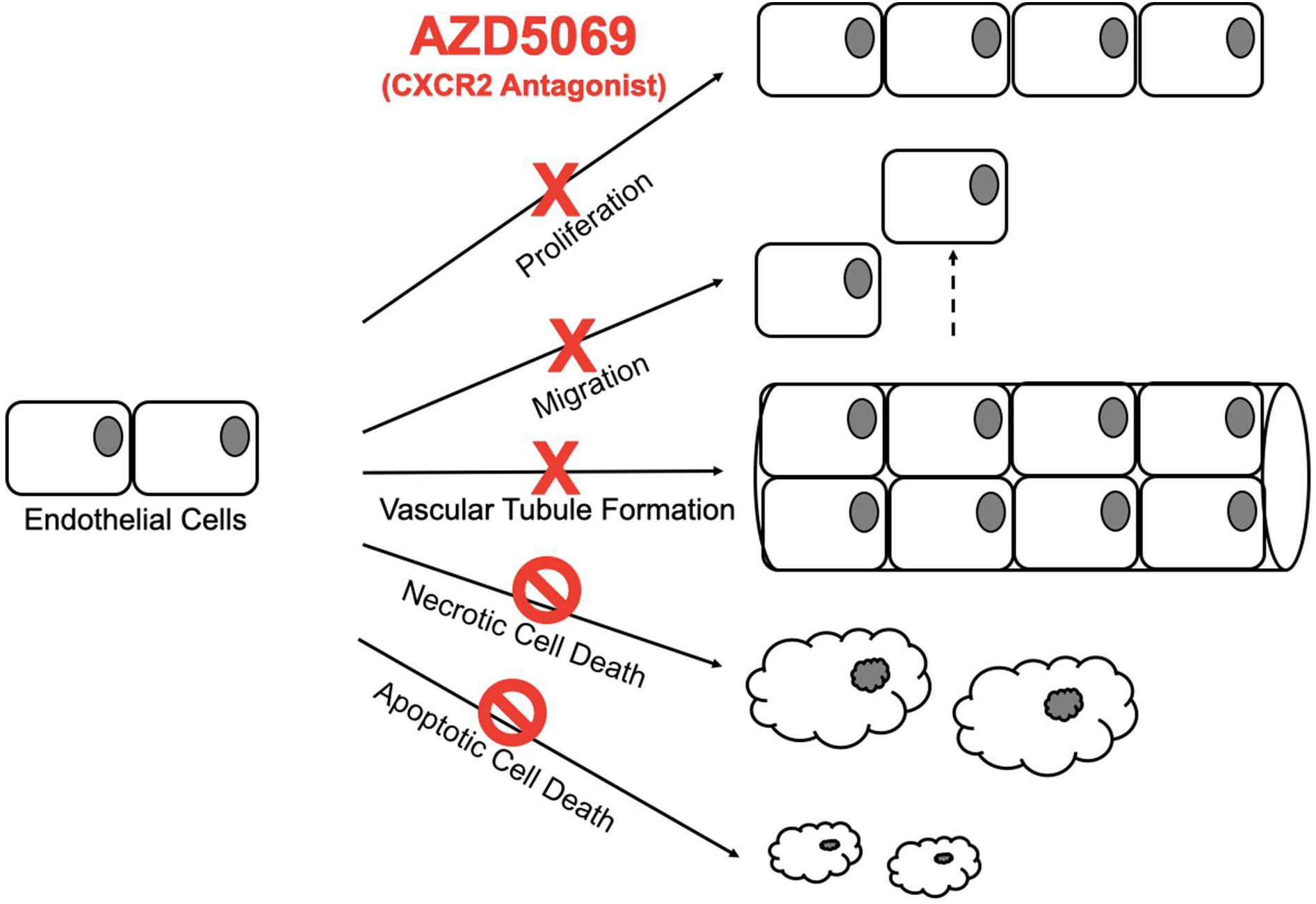

## Background

The CXCR2 receptor pathway plays a major role regulating reactive angiogenesis in inflammation and invasive angiogenesis in cancer^1–3^. CXCR2 acts as a master regulator that mobilizes bone marrow leukocytes to inflamed tissues or tumor microenvironments to orchestrate neovascularization^4,5^. In parallel, activation of CXCR2 on endothelial cells facilitates leukocyte transmigration across the blood-endothelial barrier into tissues to promote local angiogenesis^5–7^. In diseases with chronic inflammation, hyperactivation of CXCR2 may cause reactive tissue hypervascularization and sequelae^1,3,8,9^. Similarly, dysregulation of CXCR2 may be a common mechanism for tumor invasion and metastasis across multiple human cancers^2,10–15^.

The CXCR2 receptor pathway has been evaluated in humans as a target for therapy in inflammatory disease^1,3,16–18^ and cancer^1–3,11–15,19,20^. However, endothelial cell CXCR2 blockade has never been reported as an approach to inhibit pathologic angiogenesis. Ample evidence demonstrates that activation of CXCR2 on endothelial cells promotes endothelial cell proliferation, migration, and survival and increases angiogenesis^19–24^. Therefore, direct antagonism of the endothelial cell CXCR2 receptor pathway may have clinical utility to limit pathologic, proliferative, or excessive angiogenesis.

We investigated AZD5069, a clinical stage, direct CXCR2 antagonist, as a novel angiogenesis blocker *in vitro*. We tested the <u>hypothesis</u> that CXCR2 blockade with AZD5069 inhibits endothelial cell proliferation, migration, and tubule formation in human endothelial cell culture without cytotoxicity.

## Methods

Study approval was obtained from Geisinger Medical Center Institutional Review Board (IRB# 2021-0441, approved 01/05/2022) to use human blood from volunteer donors. Fresh whole blood was obtained via veni-puncture from healthy adult donors in sodium heparin blood tubes. Plasma was stored at -80 °C.

### Cell Culture Protocol

Human Umbilical Vein Endothelial Cells (HUVECs), Human Pulmonary Artery Endothelial Cells (HPAECs), Human Aortic Endothelial Cells (HAECs) (Lonza) were cultured in buffered endothelial growth media (EGM, Lonza) in T75 culture flasks at 37°C with 5% CO_2_. Fourth or fifth passage cells were used. Assays were performed with 1:1 endothelial basal media (EBM):human plasma. AZD5069 (AbCam) dilutions (0, 8, 16, 32, 64, 128, 256 μM) were mixed in EBM. Cells grown in EGM were used as a control.

Five experiments with three cell lines (HUVECs, HAECs, HPAECs) were designed to observe effects of AZD5069 on endothelial cell proliferation, migration, tubule formation, apoptosis, and necrotic cell death. Each assay was performed separately with fresh plasma from at least four human donors in duplicate. A Zeiss LSM 710 confocal microscope and Olympus phase contrast microscope were used for imaging. Image J software analyzed images.

### Endothelial Cell Proliferation

Fibronectin-coated wells were plated with 9,000 cells/cm². Proliferation was quantified by uptake of fluorescent-labeled 5-Ethynyl-2’-deoxyuridine (EdU), a nucleoside analogue, into newly synthesized DNA, as an indicator of active cell division. Nuclei were counterstained with 4′,6-diamidino-2-phenylindole (DAPI). Cell proliferation rate was reported as percentage EdU-positive nuclei relative to total nuclei per unit area as we have reported^25^. A positive control was performed with cells grown in EGM.

### Endothelial Cell Migration

Fibronectin-coated wells were plated with 30,000 cells/cm² and brought to confluence. A pipette tip created a controlled scratch across the endothelial monolayer to simulate intimal injury. Migration into the injury was quantified. Cells were imaged at 12 hours (HUVECs) or 24 hours (HPAECs, HAECs) with light microscopy to determine migration as area occupied by migrated cells relative to original scratch area as we have reported^25,26^. A positive control was performed with cells grown in EGM.

### Endothelial Cell Tubule Formation

Geltrex extracellular matrix construct was plated with 30,000 cells/cm². Endothelial cell sprouting, arborization, and formation of vascular connections were observed. Cells were imaged at 12 hours (HUVECs) or 24 hours (HPAECs, HAECs) with phase-contrast microscopy to determine endothelial tubule count, total tubule length, and node count as we have reported^25,26^. A positive control was performed with cells grown in EGM.

### Endothelial Cell Apoptosis

Fibronectin-coated wells were plated with 9,000 cells/cm². Apoptosis was quantified with *in situ* terminal transferase 2′-deoxyuridine 5′-triphosphate nick-end labeling (TUNEL) staining which catalytically incorporates fluorescein-labeled 2’-Deoxyuridine triphosphate-‘5 (dUTP) at DNA strand breaks in cells during programmed cell death. Nuclei were counterstained with DAPI. Apoptosis rate was quantified as proportion of TUNEL-positive nuclei relative to total nuclei per unit area as we have reported^25,27–30^. A positive control was performed with cells grown in EGM and DNAse to induce widespread apoptosis and TUNEL positivity.

### Endothelial Cell Necrosis

Fibronectin-coated wells were plated with 9,000 cells/cm². Cells were treated with membrane-impermeable fluorescent-labeled cyanine dye which enters cells without an intact cell membrane and incorporates into DNA to indicate necrotic cell death. Nuclei were counterstained with DAPI. Necrotic cell death was quantified as percentage green nuclei relative to total nuclei per unit area as we have reported^25,26^. A positive control was performed with cells grown in EGM and 70% methanol to induce widespread cellular lysis and dye uptake.

### Statistics

Prism software (version 9, Graph Pad Software) was used to perform statistical analyses, plot data, and build figures. Repeated measures one-way analysis of variance (ANOVA) with Tukey post hoc test was performed for each assay with each cell line. Descriptive data in the text and figures are presented as mean±standard deviation. A value of p<0.05 was considered statistically significant.

## Results

AZD5069 significantly reduced HUVEC, HAEC, and HPAEC proliferation (**Figure 1**), migration (**Figure 2**), tubule count, total tubule length, and node count (**Figure 3**) versus control. A dose-response was observed in each assay.

**Figure 1.**
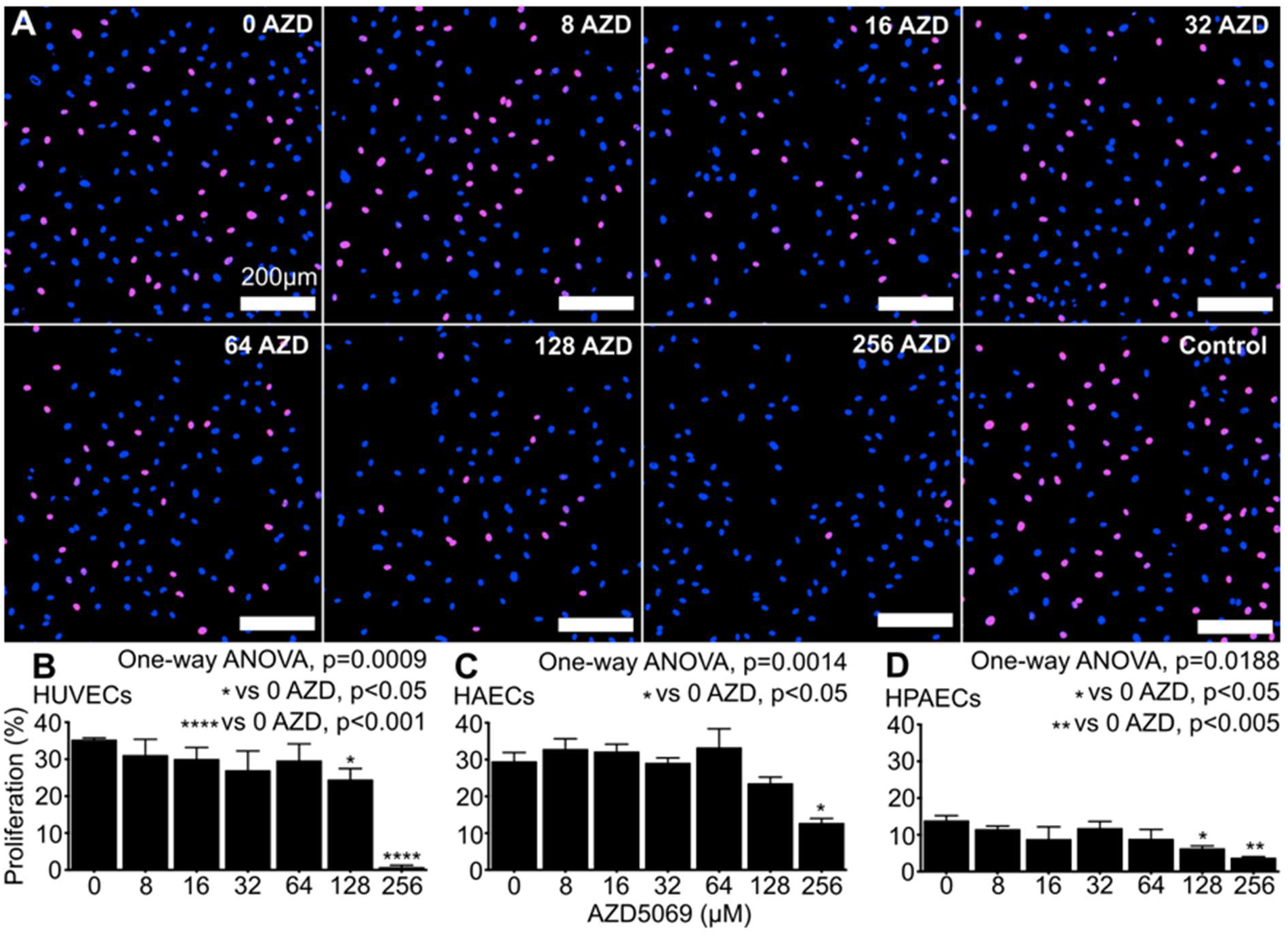
**A:** A representative experiment is shown with Human Umbilical Vein Endothelial Cells (HUVECs) grown in 1:1 human plasma:endothelial basal media across AZD5069 doses. Control, endothelial growth media. **B-D:** AZD5069 reduced HUVEC, Human Aortic Endothelial Cell (HAEC), and Human Pulmonary Artery Endothelial Cell (HPAEC) proliferation in human plasma evaluated with 5-Ethynyl-2’-deoxyuridine (EdU) uptake. AZD, AZD5069; ANOVA, analysis of variance

**Figure 2.**
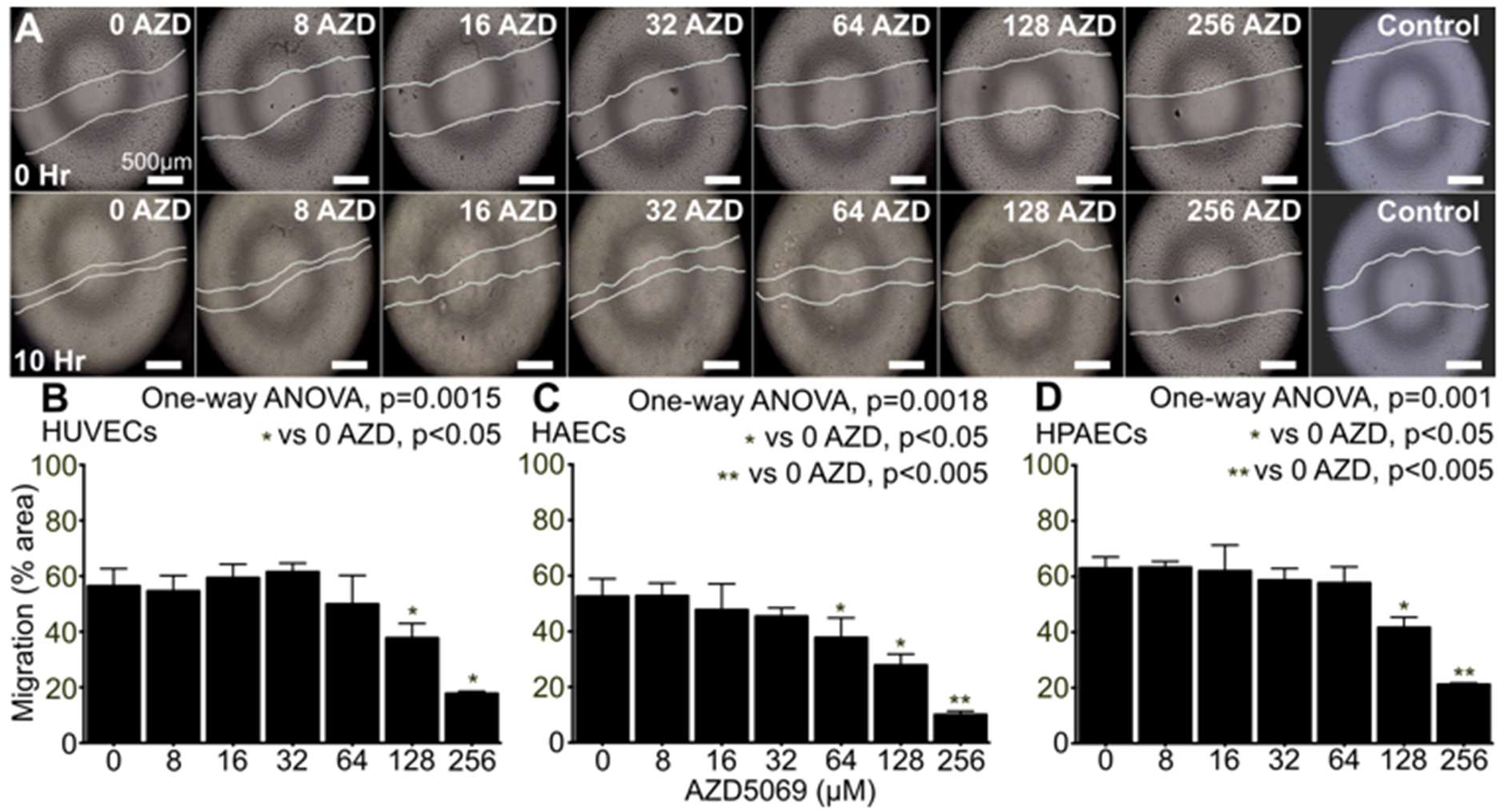
**A:** A representative experiment is shown with Human Umbilical Vein Endothelial Cells (HUVECs) grown in 1:1 human plasma:endothelial basal media across AZD5069 doses. Control, endothelial growth media. **B-D:** AZD5069 reduced HUVEC, Human Aortic Endothelial Cell (HAEC), and Human Pulmonary Artery Endothelial Cell (HPAEC) migration in human plasma evaluated with a scratch-migration assay. AZD, AZD5069; ANOVA, analysis of variance

**Figure 3.**
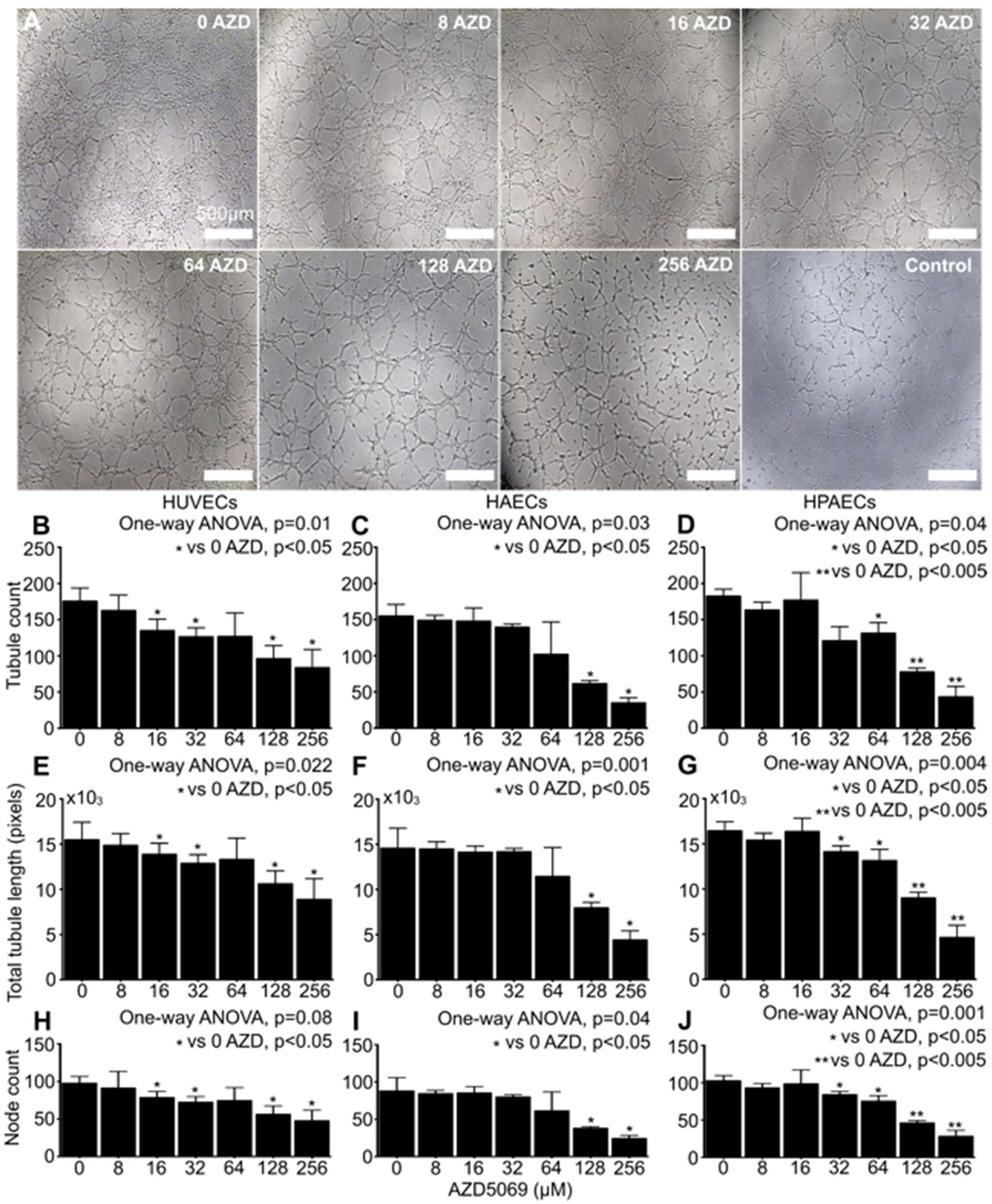
**A:** A representative experiment is shown with Human Umbilical Vein Endothelial Cells (HUVECs) grown in 1:1 human plasma:endothelial basal media across AZD5069 doses. Control, endothelial growth media. **B-D:** AZD5069 reduced HUVEC, Human Aortic Endothelial Cell (HAEC), and Human Pulmonary Artery Endothelial Cell (HPAEC) angiogenesis as determined by endothelial cell tubule count, total tubule length, and node count in human plasma evaluated by Geltrex assay. AZD, AZD5069; ANOVA, analysis of variance

AZD5069 did not cause HUVEC, HAEC, or HPAEC apoptosis (**Figure 4**) or necrotic cell death (**Figure 5**) at any concentration versus control. Cells appeared healthy without changes in confluence, orientation, or size.

**Figure 4.**
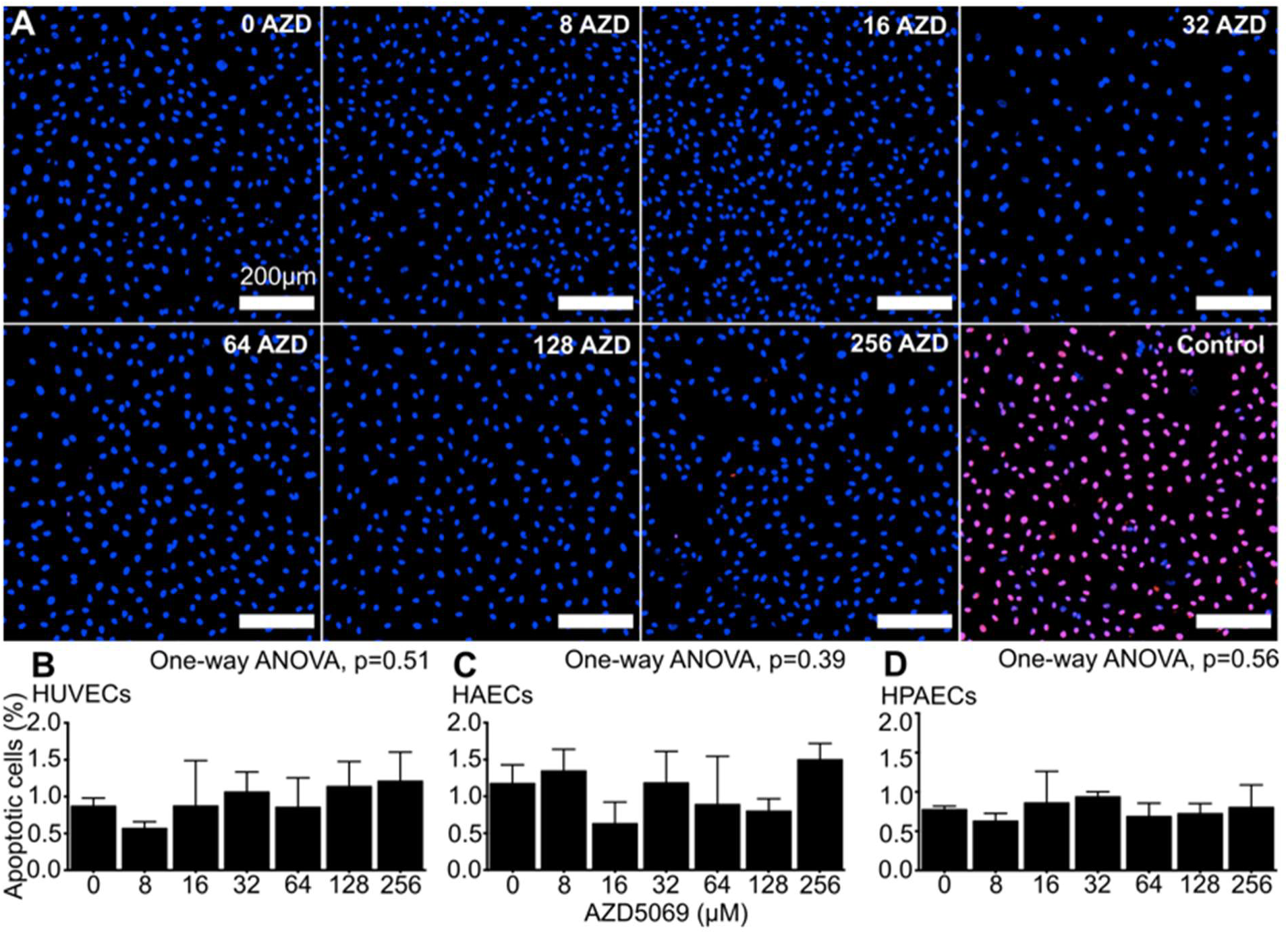
**A:** A representative experiment is shown with Human Umbilical Vein Endothelial Cells (HUVECs) grown in 1:1 human plasma:endothelial basal media across AZD5069 doses. Control, endothelial growth media and DNAse. **B-D:** AZD5069 did not cause HUVEC, Human Aortic Endothelial Cell (HAEC), or Human Pulmonary Artery Endothelial Cell (HPAEC) apoptosis quantified by *in situ* terminal 2’-Deoxyuridine triphosphate-‘5 (dUTP) nick-end labeling (TUNEL) in human plasma. AZD, AZD5069; ANOVA, analysis of variance

**Figure 5.**
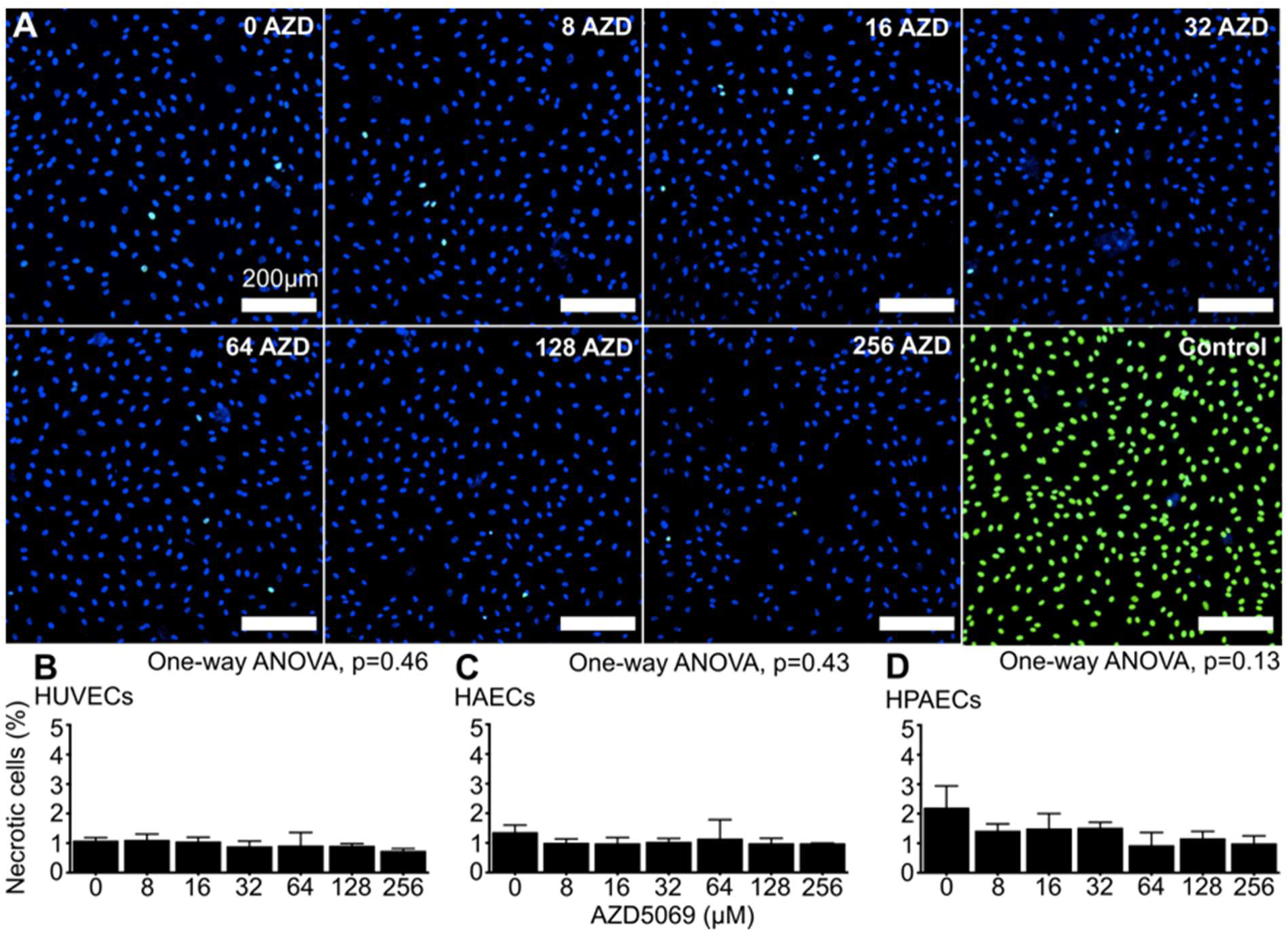
**A:** A representative experiment is shown with Human Umbilical Vein Endothelial Cells (HUVECs) grown in 1:1 human plasma:endothelial basal media across AZD5069 doses. Control, endothelial growth media and 70% methanol. **B-D:** AZD5069 did not cause HUVEC, Human Aortic Endothelial Cell (HAEC), or Human Pulmonary Artery Endothelial Cell (HPAEC) necrotic cell death in human plasma quantified by membrane-impermeable cyanine dye uptake. AZD, AZD5069; ANOVA, analysis of variance

## Discussion

We evaluated AZD5069, a selective CXCR2 antagonist, as an angiogenesis inhibitor with human endothelial cell culture. HUVECS, HAECs, and HPAECs grown with AZD5069 exhibited statistically significant dose-response reduction in endothelial cell 1) proliferation, 2) migration, and 3) tubule count, total tubule length, and node count. Cytotoxicity experiments demonstrated that AZD5069 4) did not increase apoptosis, and 5) did not increase necrotic cell death. These findings support early feasibility for endothelial cell CXCR2 blockade as an anti-angiogenesis agent. We speculate that endothelial cell CXCR2 antagonism may have utility in human diseases in which pathologic, proliferative, or excessive angiogenesis occur.

### The CXCR2 Receptor Pathway in Angiogenesis

The CXCR2 receptor pathway plays a major role regulating angiogenesis in inflammation and cancer^1–5^. CXCR2 acts as a master regulator that activates and mobilizes neutrophils and myeloid-derived suppressor cell from bone marrow into the blood stream to sites of acute and chronic inflammation, tissue injury, and tumor microenvironments. CXCR2 activated inflammatory cells enter inflamed or neoplastic tissues and orchestrate neovascularization and local tissue growth. In diseases with unregulated inflammation such as chronic obstructive airway disease (COPD), asthma, bronchiectasis, atherosclerosis, ulcerative colitis, and arthritis, unregulated CXCR2 activation may cause a reactive hypervascularization and sequelae^1,3,8,9^.

Human endothelial cells also express CXCR2 receptors that promote angiogenesis independent of inflammatory cells^20–22^. Activation of endothelial cell CXCR2 increase endothelial cell proliferation, migration, and survival^19,23,24^. These proangiogenic endothelial cell behaviors also promote a reactive angiogenesis, which if unregulated, may cause excessive growth of microvasculature.

Dysregulation of CXCR2 may be a common mechanism for invasion across multiple human cancer types^1–5^. In patients with melanoma^10^, pancreatic cancer^11^, metastatic lung cancer^12,13^, kidney cancer^20^, and breast cancer^12,14,15^, CXCR2 activation increases tumor vascularization, growth, invasion, and metastasis^10–15^. Taken together, roles of CXCR2 in inflammatory disease, reactive tissue angiogenesis, and cancer invasion suggest clinical utility for pharmacologic CXCR2 blockade, which may have applications in multiple human diseases.

### Clinical Investigation Of CXCR2 Blockade In Inflammatory Disease and Cancer

AZD5069, a clinical-stage direct CXCR2 blocker, was originally developed to inhibit neutrophils in patients with COPD^16–18^. In eight Phase I^16^ and two Phase II^17,18^ clinical trials, AZD5069 was demonstrated safe and well tolerated in humans. AZD5069 was administered to 200 healthy volunteers, and dosing, pharmacokinetics, and safety profile were reported in detail^16^. AZD5069 did significantly reduce blood neutrophil counts with a dose response. However, AZD5069 did not demonstrate clinical efficacy to relieve COPD, asthma, or bronchiectasis, and this approach was abandoned.

A therapeutic role for CXCR2 blockade as an immuno-modulator of inflammatory cells is also being investigated in cancer. Multiple CXCR2 inhibitors are under investigation. SB225002, SCH-527123, SCH-479833, Ladarixin (DF2156A), Reparixin, monoclonal antibody anti-human CXCL1 (HL2401), and AZD5069 are each speculated to have possible utility in cancer therapeutics^2,11,12,15,19^. Specifically, AZD5069 is being evaluated as an adjunct to traditional chemo therapy in multiple clinical trials (ClinicalTrials.gov ID: NCT02583477; NCT02499328; NCT03177187; ISRCTN12669009). These trials aim to “prime” the immune system by inhibiting CXCR2-dependant neutrophil recruitment in the tumor microenvironment.

### CXCR2 Blockade To Treat Proliferative Angiogenesis

CXCR2 blockade with AZD5069 has not been investigated nor reported as an anti-angiogenesis agent of which we are aware. We have demonstrated with multiple human endothelial cell lineages that AZD5069 decreased endothelial cell proliferation, migration, and tubule formation. As expected, cytotoxic effects were not observed. This is consistent with the favorable safety profile of AZD5069 reported in human clinical trials^16–18^. As such, we speculate that AZD5069 may have clinical utility to limit proliferative angiogenesis in diseases with pathologic, dysregulated, or excessive angiogenesis. Pharmacologic blockade to limit endothelial cells from dividing, migrating, and forming new vascular tubules, which are inciting steps of proliferative angiogenesis, is conceptually appealing, especially if accomplished without causing endothelial cell death. Many clinical examples of proliferative angiogenesis exist. Angiodysplasia, telangiectasia, vascular ectasia, collateral vascular channel growth, and arteriovenous malformations are collectively proliferative vascular lesions characterized by disorganized overgrowth of vessels with abnormal vascular architecture and function. These lesions commonly present in patients with single-ventricle congenital heart disease^25,31–35^, hereditary hemorrhagic telangiectasia^36^, gastric antral vascular ectasia^37^, Heyde’s syndrome^38,39^, congenital and acquired von Willebrand syndrome^38,40–42^, hepatopulmonary syndrome^43^, non-pulsatile ventricular assist devices^26,44,45^, and elderly patients^46^. Limited therapy exists to prevent or treat these lesions. Pharmacologic blockade of angiogenesis without cytotoxic effects to endothelial cells has obvious applications and utility in medicine and biology. We are performing ongoing investigations to identify specific pathologies and patient populations that may benefit from CXCR2 blockade with AZD5069 therapy.

### Study Limitations

This study was conducted with human plasma and human cells *in vitro*. Cell culture does not replicate complex and dynamic 3-dimensional environments of a living organism and may poorly predict human responses and efficacy *in vivo*.

## Conclusions

AZD5069 inhibited human umbilical, aortic, and pulmonary endothelial cell proliferation, migration, and tubule formation without cytotoxicity in human endothelial cell culture. Blockade of the endothelial cell CXCR2 receptor pathway may have clinical utility to treat pathologic, dysregulated, or excessive angiogenesis in cardiovascular, immunologic, inflammatory, and oncologic disease. Further investigation is warranted.

## Acknowledgments

Support for this project was provided by NIH R01 HL172022, NIH R01 HL166334, and the Janet Weis Children’s Hospital Beyond the Bricks Foundation. We acknowledge and thank Samson Hennessy-Strahs, Damian Mason, and Stephanie Buczkowski for assistance and support.

## Clinical Perspectives

The CXCR2 receptor pathway plays a major role in inflammatory and invasive angiogenesis. We evaluated AZD5069, a clinical-stage direct CXCR2 antagonist, as a novel angiogenesis inhibitor. In human cell culture, AZD5069 inhibited angiogenesis without cytotoxicity. The endothelial cell CXCR2 receptor pathway may be a novel target for anti-angiogenesis therapy. AZD5069 may have clinical utility as a novel angiogenesis blocker in cardiovascular, inflammatory, and oncologic disease.

## Translational Outlook

AZD5069 inhibited endothelial cell proliferation, migration, and tubule formation without causing apoptosis or necrotic cell death in human endothelial cell culture. Pharmacologic antagonism of the endothelial cell CXCR2 receptor pathway may have clinical utility to prevent or treat pathologic, dysregulated, or excessive angiogenesis in human disease. Further investigation is warranted to identify specific human pathologies that may benefit from CXCR2 blockade.

## Highlights

- The CXCR2 receptor pathway plays a major role regulating angiogenesis in inflammation and cancer.
- The CXCR2 receptor pathway has been evaluated in humans as a target for therapy in inflammatory disease and cancer but not as a therapeutic approach to block pathologic angiogenesis.
- AZD5069 is a clinical stage, direct CXCR2 antagonist.
- In human endothelial cell culture, AZD5069 inhibited angiogenesis without causing apoptosis or necrotic cell death.
- The endothelial cell CXCR2 receptor pathway may be a novel target for anti-angiogenesis therapy.
- AZD5069 may have clinical utility as a novel angiogenesis blocker in human disease.

## Abbreviations

EdU: 5-Ethynyl-2’-deoxyuridine
dUTP: 2’-Deoxyuridine triphosphate-‘5
EGM: endothelial cell growth media
EBM: endothelial cell basal media
HUVEC: Human Umbilical Venous Endothelial Cell
HAEC: Human Aortic Endothelial Cell
HPAEC: Human Pulmonary Artery Endothelial Cell
DAPI: 4′,6-diamidino-2-phenylindole
TUNEL: terminal deoxynucleotidyl transferase 2’-Deoxyuridine triphosphate-‘5 nick-end labeling
ANOVA: analysis of variance

